# Differential quantitative requirements for pre-mRNA splicing-regulated shelterin protein levels in distinct telomere functions

**DOI:** 10.1101/2025.09.02.673742

**Authors:** Miho Takeuchi, Yoko Otsubo, Junko Kanoh

**Affiliations:** Former address: Institute for Protein Research, Osaka University, 3-2 Yamadaoka, Suita, Osaka 565-0871, Japan; Life Science Network, the University of Tokyo, Komaba 3-8-1, Meguro-ku, Tokyo 153-8902, Japan; Department of Life Sciences, Graduate School of Arts and Sciences, the University of Tokyo, Komaba 3-8-1, Meguro-ku, Tokyo 153-8902, Japan; Department of Biological Sciences, Graduate School of Science, the University of Tokyo, Hongo 7-3-1, Bunkyo-ku, Tokyo 113-0033, Japan; Collaborative Research Institute for Innovative Microbiology (CRIIM), the University of Tokyo, Yayoi 1-1-1, Bunkyo-ku, Tokyo 113-0032, Japan

## Abstract

Telomeres perform multiple functions to maintain genome stability, including telomere length regulation, chromosome end protection, and meiotic chromosome dynamics. These functions are governed by shelterin, a telomere-binding protein complex. Here, we show that efficient pre-mRNA splicing of the *Schizosaccharomyces pombe* shelterin components Rap1 and Poz1 ensures sufficient protein levels, which are critical for telomere maintenance. Our analyses revealed that Tls1 and Cay1 act at distinct steps in splicing, specifically affecting *rap1*⁺ and *poz1*⁺ transcripts: Tls1 strongly interacts with Brr2 (a splicing factor), whereas Cay1 preferentially associates with introns. Accordingly, deletion of *tls1*⁺ and *cay1*⁺ synergistically impaired splicing of *rap1*⁺ and *poz1*⁺ transcripts and reduced their protein levels, leading to abnormal telomere elongation. Removal of introns from the *rap1*⁺ and *poz1*⁺ genes restored normal protein levels and telomere length, confirming that defective splicing underlies these defects. Analyses of the phenotypes of single and double *tls1*Δ and *cay1*Δ mutants revealed that different telomere functions vary in their dependence on Rap1 levels: telomere length regulation and, to a lesser extent, meiosis require higher protein abundance, whereas chromosome end protection can be sustained with minimal amounts. These findings reveal a hierarchical requirement for Rap1 across telomere functions and establish a framework for understanding how splicing-dependent regulation of shelterin components coordinates multiple aspects of telomere biology.

## INTRODUCTION

Telomeres are specialized nucleoprotein structures located at the ends of linear chromosomes. Due to the end-replication problem and the susceptibility of chromosome termini to be recognized as DNA double-strand breaks, eukaryotic cells have evolved telomeres to prevent genome instability. Telomeric DNA consists of tandemly repeated, species-specific sequences, which are bound by dedicated telomere-binding proteins. These proteins assemble into a higher-order structure known as the shelterin complex, which physically bridges the single-stranded (ss) overhang at the extreme chromosome terminus with the adjacent double-stranded (ds) telomeric region (1,2). Shelterin plays dual roles: it protects chromosome ends from inappropriate DNA repair activities, such as end-to-end fusions, and regulates telomerase recruitment to maintain telomere length homeostasis (1,2).

In the fission yeast *Schizosaccharomyces pombe* (*S. pombe*), the shelterin complex is composed of six subunits: Taz1, Rap1, Poz1, Ccq1, Tpz1, and Pot1 (3). Among them, Taz1—a functional homolog of mammalian TRF1 and TRF2—directly binds to telomeric dsDNA, while Pot1—a homolog of mammalian POT1—associates with telomeric ssDNA (4,5). These two DNA-binding factors are interconnected through a protein-protein interaction network: Taz1 interacts with Rap1 (a homolog of mammalian RAP1 and *Saccharomyces cerevisiae* [*S. cerevisiae*] Rap1), which associates with Poz1, which in turn binds to Tpz1 associated with Ccq1 and Pot1 (3,6,7) (Figure 1A).

**Figure 1.**
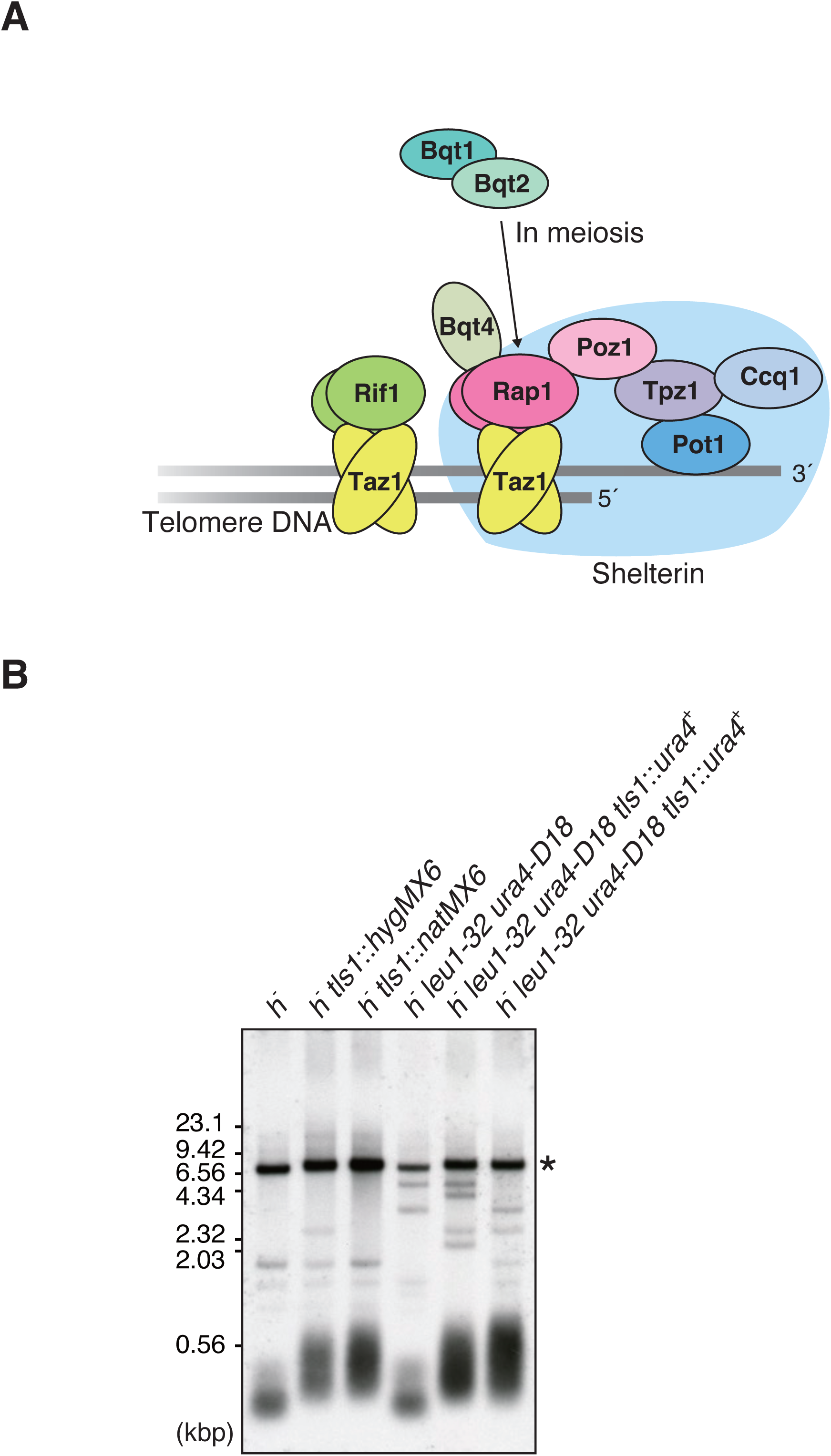
Deletion of the *tls1* gene results in moderate telomere elongation. (A) Telomere-binding proteins in *S. pombe*. The shelerin complex (shaded in pale blue) bridges ssDNA and dsDNA at telomeres to protect and maintain telomere DNA. Taz1 associates with Rif1, a global regulator of DNA replication timing. Rap1 interacts with Bqt4 to tether telomeres to the nuclear envelope and with Bqt1 and Bqt2 to mediate telomere clustering during meiosis. (B) Analysis of telomere length in the wild-type and *tls1Δ* strains in various genetic backgrounds. Genomic DNA was digested with ApaI, separated in a 1% agarose gel, and analyzed by Southern blotting using telomeric DNA as a probe. Smeared signals at the bottom represent telomeres of chromosomes 1 and 2, and an asterisk indicates the position of chromosome 3 telomeres.

Rap1 functions as a central scaffold within shelterin, mediating multiple telomere-associated processes (8). Through interaction with Poz1, Rap1 contributes to negative regulation of telomere length (3). Additionally, Rap1 connects telomeres to the nuclear envelope via the nuclear membrane protein Bqt4 and facilitates chromosome movements during mitosis and meiosis (9,10). The Taz1-Rap1-Poz1 axis is essential for maintaining telomere length within a defined range (approximately 300 bp in *S. pombe*), while the Tpz1-Ccq1-Pot1 module is required for recruitment of telomerase, whose catalytic subunit Trt1 extends telomeric DNA and ensures chromosome end protection (3). Furthermore, Taz1 interacts with Rif1, a key factor that coordinates telomere length regulation and the temporal control of DNA replication at late-firing origins across the genome (7,11,12) (Figure 1A).

In this study, we reveal a regulatory mechanism governing the expression of two shelterin components, Rap1 and Poz1, at the post-transcriptional level. Specifically, we show that *rap1*^+^ and *poz1*^+^ mRNA expression is modulated through a shared pathway involving pre-mRNA splicing. Through a yeast two-hybrid screen in *S. cerevisiae*, we identified Rip6 (<u>R</u>ap1-interacting <u>p</u>rotein 6), also known as Tls1 (13), as a candidate Rap1-interacting protein. Unexpectedly, in *S. pombe*, Rip6/Tls1 does not exhibit direct interaction with Rap1 *in vivo*; rather, it regulates the splicing of *rap1*^+^ and *poz1*^+^ transcripts in cooperation with Cay1, a previously identified splicing factor for these genes (14,15). Our findings demonstrate that proper pre-mRNA splicing of *rap1*^+^ and *poz1*^+^ is required to produce adequate levels of Rap1 and Poz1 proteins, which in turn are essential for normal telomere functions. We further reveal that different telomere functions vary in their quantitative requirements for Rap1: telomere length regulation, and to a lesser extent meiosis, require higher protein levels, whereas chromosome end protection can be maintained with only minimal amounts. Thus, this study uncovers a hierarchical requirement for Rap1 within distinct telomere functions, providing new insight into the post-transcriptional regulation of shelterin-mediated telomere control.

## MATERIALS AND METHODS

### Strains, media and general techniques for *S. pombe*

The *S. pombe* strains used in this study are listed in Supplementary Table S1. Growth media and basic genetic and biochemical techniques were described previously (16–18).

### Screen for genes encoding Rap1-interacting proteins

The entire *rap1*^+^ ORF was cloned into pGBKT7 (Clontech) to produce pGBKT7-*rap1*^+^. The *S. cerevisiae* strain Y190 (*MATa ura3-52 his3-200 lys2-801 ade2-101 trp1-901 leu2-3 112 gal4*Δ *gal80*Δ *URA3*::*GAL1_UAS_-GAL1_TATA_*-*lacZ cyh^r^2 LYS2*::*GAL1_UAS_-HIS3_TATA_-HIS3 MEL1*) was co-transformed with pGBKT7-*rap1*^+^ and *S. pombe* MATCHMAKER cDNA library constructed in pGAD-GH (Clontech). Approximately 574 million transformants were screened for *HIS3* expression on SD media lacking histidine but containing 25 or 30 mM 3-AT (3-Amino-1,2,4-triazole), a competitive inhibitor of *HIS3*. A total of 500 His^+^ colonies were obtained. 200 transformants showing high β-galactosidase activity were selected, and the cDNAs in pGAD-GH were isolated from these transformants using *Escherichia coli* (*E. coli*, DH5α). The Y190 strain was then re-transformed with pGBKT7-*rap1*^+^ and each isolated cDNA in pGAD-GH, and β-galactosidase activity was assayed. Finally, 63 transformants were confirmed to be positive for β-galactosidase activity, and the cDNAs were sequenced.

### Yeast two-hybrid assay

The entire ORFs of various genes were individually cloned into pACT2 (Clontech) or pGBKT7 (Clontech). The *S. cerevisiae* strain Y190 was used for β-galactosidase activity assays.

### Production and purification of anti-Tls1 antibody

To produce polyclonal antibodies against Tls1, the whole coding region of the *tls1* gene was cloned into pGEX-5X-2 (Cytiva). The GST-Tls1 protein was purified from *E. coli* BL21-CodonPlus (Stratagene) transformed with pGEX-5X-2-*tls1*^+^ and used to immunize rabbits. The antibodies were affinity purified using GST-Tls1 protein on Immobilon-P transfer membranes (Merck Millipore).

### Southern blotting

To analyze telomere DNA lengths, genomic DNA (20 μg) was digested with ApaI, separated in 1% agarose gel, and transferred to Hybond-N^+^ nylon membranes (GE Healthcare Life Science). To generate the telomere probe, the telomeric DNA fragment was labelled with digoxigenin using the DIG High Prime DNA Labelling and Detection Starter Kit II (Roche).

### Western blotting

To analyze protein abundance, exponentially growing cells were collected and treated with 150-200 µl of lysis buffer (1.85 M NaOH, 7.4% 2-mercaptoethanol) for 10 min and mixed with the same volume of 50% trichloroacetic acid (TCA) for 10 min on ice. After centrifugation at 14,000 rpm for 2 min, pellets were denatured in SDS-PAGE loading buffer and analyzed in 8% SDS-PAGE gels. The whole cell extracts were transferred to Immobilon-P Transfer Membrane (Millipore) and probed with anti-Rap1 (10), anti-Flag (M2 F3165, Sigma), and anti-PSTAIR (P7962, Sigma) antibodies to detect Rap1, Flag-tagged proteins, and Cdc2 (the loading control), respectively.

### RNA analyses

Total RNA was prepared from exponentially growing cells and treated with recombinant DNase I (RNase-free) (TaKaRa Bio). For reverse transcription (RT)-PCR, cDNA was synthesized using a High-Capacity cDNA Reverse Transcription Kit (Applied Biosystems) with random primers. RNA levels were analyzed by quantitative PCR (qPCR) using the StepOne^TM^ real-time PCR system (Thermo). Sequences of the primer sets used for qPCR are listed in Supplementary Table S2. For detection of splicing patterns, cDNA was amplified by PCR using the primer sets listed in Supplementary Table S2, and the PCR products were analyzed by acrylamide gel electrophoresis. The intensity of each band was quantified using ImageJ (version 1.52).

### GST pull-down assay

Full length *tls1*^+^ or *cay1*^+^ coding regions were cloned into pGEX-5X-2 (GE Healthcare). *Escherichia coli* BL21-CodonPlus (Stratagene) was transformed with the plasmid, and glutathione *S*-transferase (GST), GST-Tls1, or GST-Cay1 proteins were purified using Glutathione Sepharose 4B (GE Healthcare Life Science) in TNE buffer (40 mM Tris-HCl [pH 7.5], 150 mM NaCl, 5 mM EDTA, 1% Triton X-100, 50 mM NaF, 20 mM β-glycerophosphate). Each protein bound to glutathione beads was mixed with *S. pombe* cell extracts in IP buffer (50 mM HEPES-KOH [pH 7.5], 100 mM NaCl, 1 mM EDTA [pH 8.0], 0.5% Triton X-100, 0.1 mM Na_3_VO_4_, 50 mM NaF, 20 mM β-glycerophosphate) at 4°C for 1.5 h and washed four times with IP buffer. The protein complexes were boiled in SDS sample buffer and analyzed by SDS-PAGE, followed by western blotting and Coomassie Brilliant Blue (CBB) gel staining.

### *In vitro* RNA-protein binding assay

GST proteins bound to glutathione beads were treated with HB buffer (25 mM MOPS [pH 7.2], 50 mM NaCl, 15 mM EGTA, 15 mM MgCl_2_, 0.1 mM NaVO_3_, 60 mM β-glycerophosphate, 1 mM DTT, 1% NP-40) supplemented with 1 M NaCl at 4°C for 15 min and washed three times with HB buffer. The beads were mixed with *S. pombe* cell extracts in HB buffer at 4°C for 1.5 h and washed four times with HB buffer. RNAs bound to the beads were sequentially eluted with 30 µl, 30 µl, and 40 µl of HB buffer supplemented with 1 M NaCl. RNAs from the total 100 µl elute were precipitated with ethanol and glycogen, followed by DNase treatment, phenol extraction, and ethanol precipitation. RNA abundance was analyzed by RT-qPCR using the StepOne^TM^ real-time PCR system (Thermo).

### Pulsed-field gel electrophoresis (PFGE)

PFGE of NotI-digested chromosomal DNA was performed using a CHEF-DR III Pulsed Field Electrophoresis Systems (Bio-Rad) under the following conditions: 1% SeaKem Gold Agarose (Lonza) in 0.5 × TBE; temperature, 10°C; initial switch time, 40 s; final switch time, 80 s; run time, 18 h; voltage gradient, 6.8 V/cm; and angle, 120°. The chromosome DNA was analyzed by Southern blotting.

### Sporulation assay

Homothallic *h^90^* cells were grown on YES media at 32°C and then transferred onto MEA media at 28°C for 2 days. Cells with spores were suspended in H_2_O and observed with the microscope, Olympus BX53 (Olympus).

## Results

### Identification of the *rip6*^+^/*tls1*^+^ gene

To elucidate the unknown regulatory mechanisms governing shelterin function, we performed a yeast two-hybrid screen using *S. cerevisiae* to identify candidate Rap1-binding proteins (see Materials and Methods). Among the candidates, we identified a protein designated Rip6 (<u>R</u>ap1-interacting <u>p</u>rotein 6), which was subsequently found to be identical to Tls1 (13) (Supplementary Figure S1). Accordingly, we hereafter refer to this protein as Tls1. Although Tls1 was isolated as a Rap1-associated protein in our two-hybrid assay using *S. cerevisiae*, co-immunoprecipitation experiments using anti-Tls1 and anti-Rap1 antibodies failed to detect their interaction in *S. pombe* (data not shown).

### Moderate telomere DNA elongation in *tls1*Δ cells

To investigate the function of Tls1, we deleted the entire coding region of the *tls1*^+^ gene. The resulting *tls1*Δ strain exhibited moderately elongated telomere DNA (∼400 bp) compared with the wild type (∼300 bp) (Figure 1B), indicating that Tls1 contributes to the regulation of telomere DNA length. To confirm this observation, we independently generated multiple *tls1*Δ strains by replacing the entire *tls1*^+^ coding region with different selectable markers (*hphMX6*, *natMX6*, and *ura4*^+^) and consistently observed only moderate telomere elongation (Figure 1B). These results indicate that Tls1 exerts a relatively modest effect on telomere DNA length.

### Reduction of Rap1 and Poz1 protein levels in *tls1*Δ cells

To further elucidate the role of Tls1 in telomere length control, we examined whether the abundance of shelterin components is affected by deletion of *tls1*^+^. Whole-cell extracts from wild-type and *tls1*Δ strains were analyzed by western blotting. Notably, the protein levels of Rap1 and Poz1 were markedly reduced in *tls1*Δ cells, whereas those of Taz1, Pot1, Tpz1, Ccq1, Rif1, and Sgo2 (a shugoshin family protein, used here as a negative control) remained unchanged (Figure 2, lanes 1 and 2). These results suggest that Tls1 is selectively required for the proper expression of *rap1*^+^ and *poz1*^+^, or for the stability of their gene products.

**Figure 2.**
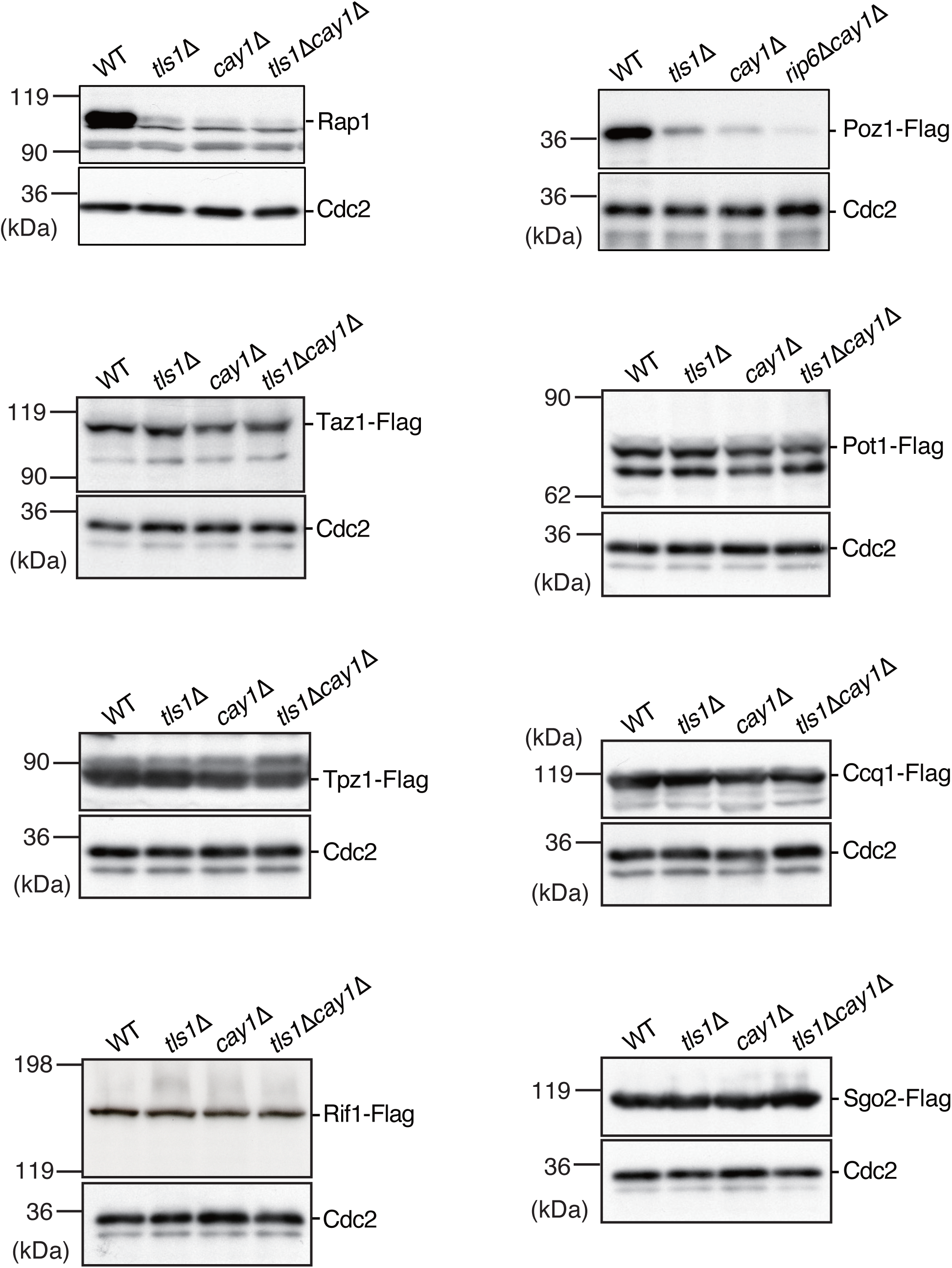
Deletion of Tls1 and Cay1 causes a severe decrease in Rap1 and Poz1 protein levels. Whole-cell extracts from wild-type (WT), *tls1*Δ, *cay1*Δ, and *tls1*Δ*cay1*Δ strains were analyzed by western blotting with anti-Rap1, anti-Flag, or anti-PSTAIRE (for Cdc2 as a loading control) antibodies.

### Tls1 controls pre-mRNA splicing of *rap1*^+^ and *poz1*^+^

To investigate whether Tls1 regulates RNA expression of *rap1*^+^ and *poz1*^+^, we quantified both exon and intron levels of their transcripts by RT-qPCR. In *tls1*Δ cells, intron levels of *rap1*^+^ and *poz1*^+^ were significantly elevated compared with those in wild-type cells, whereas no such increase was detected for the *sgo2*^+^ intron, which served as a negative control (Figure 3A and B). These results suggest that the introns of *rap1*^+^ and *poz1*^+^ are not properly spliced out in *tls1*Δ cells.

**Figure 3.**
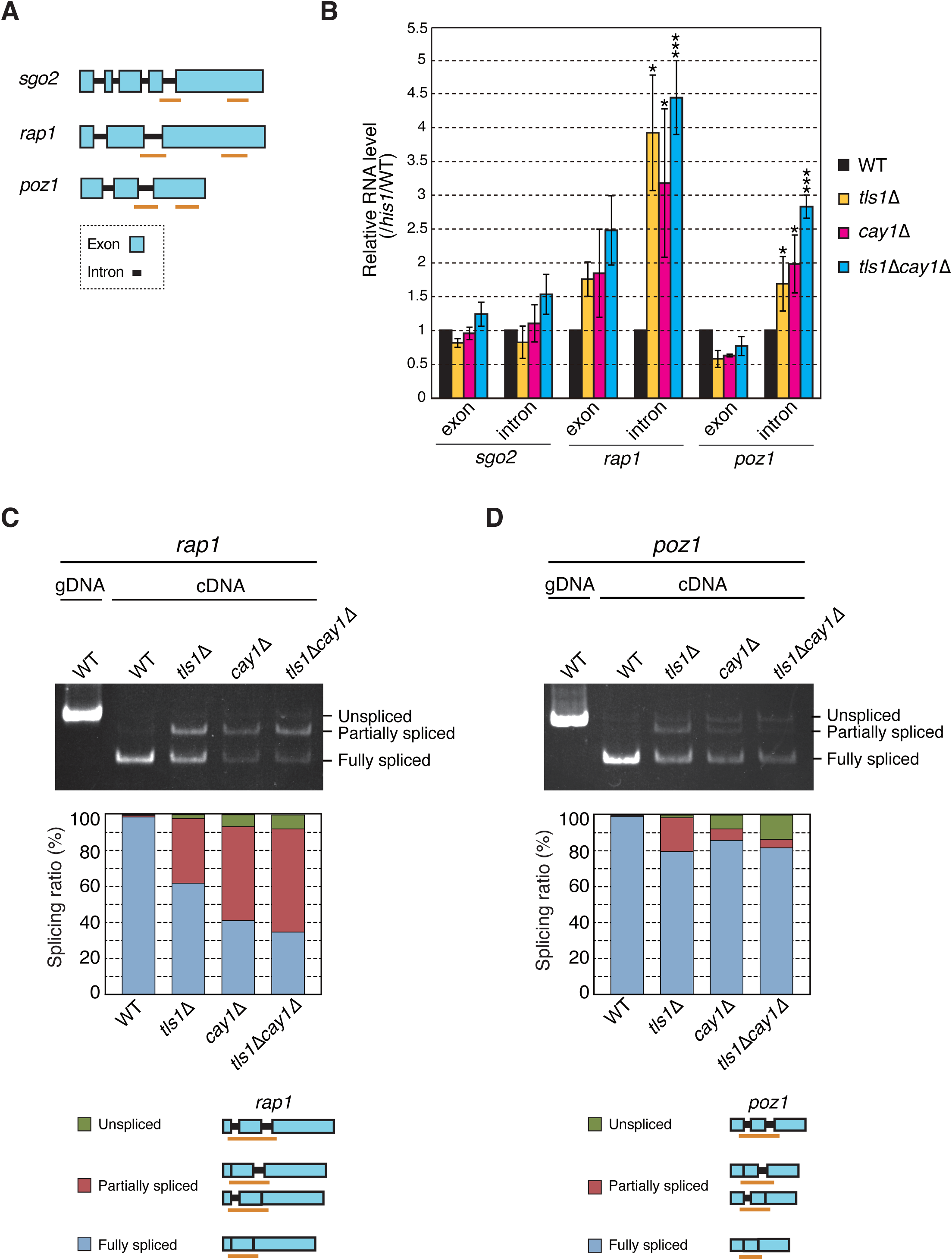
Tls1 and Cay1 control pre-mRNA splicing of *rap1*^+^ and *poz1*^+^. (A) Schematic illustration of the exon-intron structures within the coding regions of *sgo2*^+^, *rap1*^+^, and *poz1*^+^. Pale blue boxes represent exons, and black lines indicate introns. Orange bars show the DNA regions amplified by RT-qPCR in (B). (B) Levels of exonic or intronic sequences of *sgo2*^+^ (control), *rap1*^+^, and *poz1*^+^ in wild-type (WT), *tls1*Δ, *cay1*Δ, and *tls1*Δ*cay1*Δ strains were analyzed by RT-qPCR. Each value was normalized first to that of *his1*^+^ (exon) and then to the wild-type value. Error bars represent the standard deviation (s.d.) (*N* = 3 biologically independent experiments). Student’s t-test was performed against the wild type. \*\*\**p* ≤ 0.001; 0.01 < \**p* ≤ 0.05. (C) Deletion of Tls1 and Cay1 causes accumulation of unspliced or partially spliced *rap1*^+^ pre-mRNA. cDNA was synthesized from total RNA of each strain by RT-PCR and analyzed by acrylamide gel electrophoresis. Orange bars indicate the DNA regions amplified by RT-PCR using *rap1*-specific primers. Band intensities were quantified using ImageJ (version 1.52). gDNA indicates the PCR product amplified from genomic DNA with the same primers. (D) Deletion of Tls1 and Cay1 causes accumulation of unspliced or partially spliced *poz1*^+^ pre-mRNA. cDNA was analyzed as described in (C) using *poz1*-specific primers.

To directly assess the role of Tls1 in pre-mRNA splicing of *rap1*^+^ and *poz1*^+^, we analyzed splicing status of their transcripts by RT-PCR followed by gel electrophoresis. In *tls1*Δ cells, approximately 40% of *rap1*^+^ and 20% of *poz1*^+^ transcripts were incompletely spliced, whereas nearly all transcripts were fully spliced in wild-type cells (Figure 3C and D). These findings indicate that Tls1 is required for efficient splicing of *rap1*^+^ and *poz1*^+^ transcripts.

### Synergistic effects of Tls1 and Cay1 deletions on Rap1 and Poz1 protein levels

Previous studies have reported that Cay1, a spliceosome subunit, also regulates the splicing of *rap1*^+^ and *poz1*^+^ transcripts (14,15). To investigate the functional relationship between Tls1 and Cay1, we analyzed the phenotypes of the *tls1*Δ*cay1*Δ double mutant. The protein levels of Rap1 and Poz1 were reduced in *cay1*Δ cells to a greater extent than that in *tls1*Δ cells, and the double mutant *tls1*Δ*cay1*Δ exhibited even more pronounced reductions compared with either single mutant (Figure 2). Consistent with these results, *cay1*Δ and *tls1*Δ*cay1*Δ cells displayed significantly increased intron levels of the *rap1*^+^ and *poz1*^+^, along with severe splicing defects in splicing in these transcripts (Figure 3B-D). These findings suggest that Tls1 and Cay1 regulate the splicing of *rap1*^+^ and *poz1*^+^ in at least partially independent pathways.

### Tls1 and Cay1 regulate Rap1 and Poz1 protein levels via pre-mRNA splicing

To assess the contribution of *rap1*^+^ and *poz1*^+^ splicing, mediated by Tls1 and Cay1, to the steady-state protein levels of Rap1 and Poz1, we removed the intron sequences from the chromosomal *rap1*^+^ and *poz1*^+^ loci, thereby eliminating the requirement for pre-mRNA splicing of these genes. The intron-less *rap1*^+^ and *poz1*^+^ constructs produced Rap1 and Poz1 proteins at levels comparable to those observed in wild-type cells, and deletion of *tls1*^+^, *cay1*^+^, or both no longer resulted in a reduction of Rap1 or Poz1 protein levels (Figure 4A and B). Taken together with the data in Figures 2 and 3, these results demonstrate that Tls1 and Cay1 redundantly maintain Rap1 and Poz1 protein levels through the regulation of pre-mRNA splicing, with Cay1 having a greater impact than Tls1.

**Figure 4.**
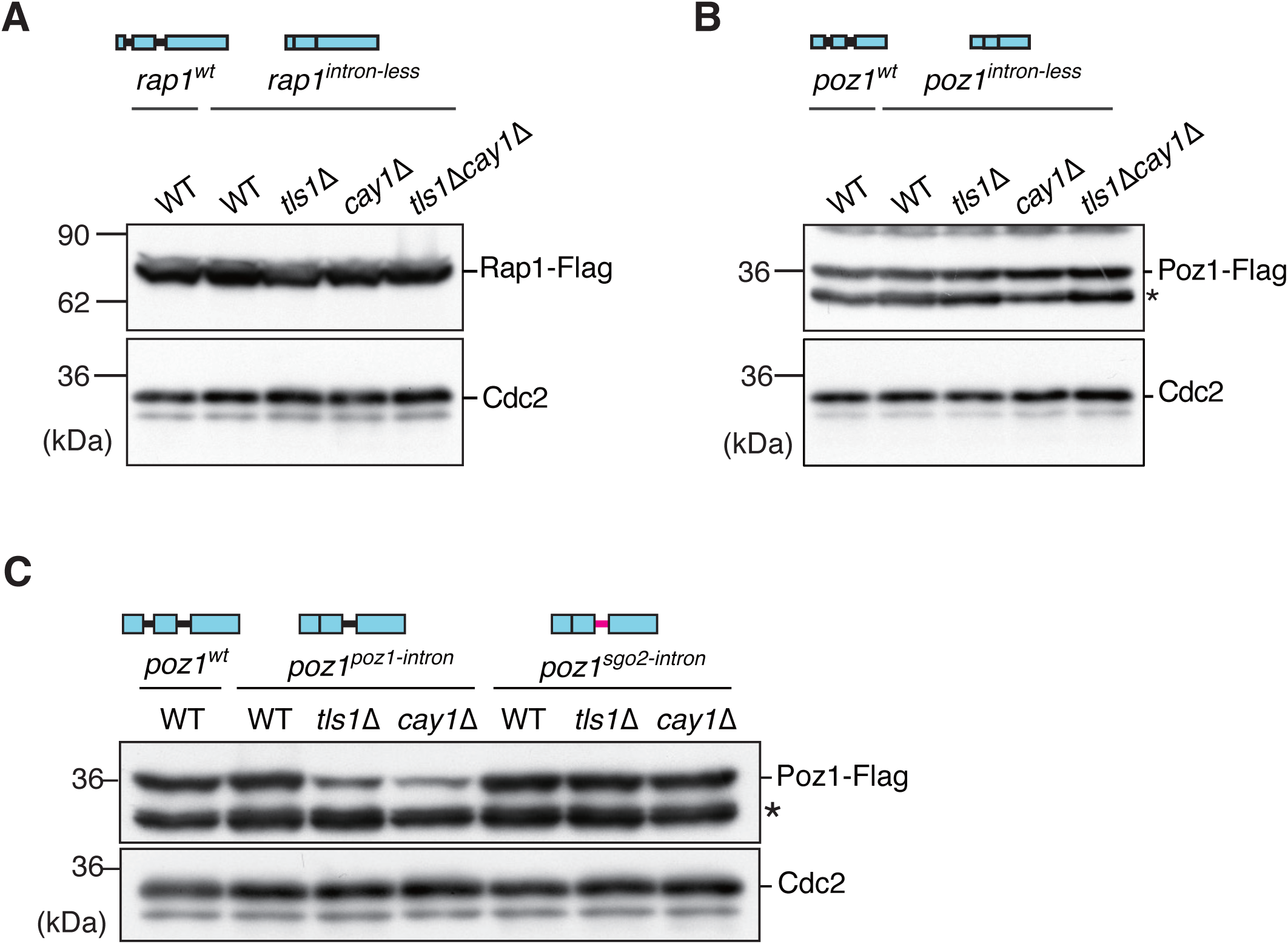
Intron-less Rap1 and Poz1 show no decrease in protein levels upon deletion of Tls1 and Cay1. (A) Whole-cell extracts from strains expressing *rap1^wt^* or *rap1^intron-less^* were analyzed by western blotting with anti-Flag (Rap1) or anti-PSTAIRE (Cdc2, loading control) antibodies. (B) Whole-cell extracts from strains expressing *poz1^wt^* or *poz1^intron-less^* were analyzed by western blotting with anti-Flag (Poz1) or anti-PSTAIRE (Cdc2, loading control) antibodies. An asterisk indicates a non-specific band. (C) Whole-cell extracts from various strains expressing *poz1^wt^*, *poz1^poz1-intron^*, or *poz1^sgo2-intron^* were analyzed by western blotting with anti-Flag (Poz1) or anti-PSTAIRE (for Cdc2, loading control) antibodies. An asterisk indicates a non-specific band.

### Intron-dependent specificity of Tls1- and Cay1-mediated splicing regulation

To examine the role of intron sequences in the specific regulation of *rap1*^+^ and *poz1*^+^ transcripts by Tls1 and Cay1, we replaced the second intron of *rap1*^+^ (83 bp) and *poz1*^+^ (94 bp) with the fourth intron of *sgo2*^+^ (52 bp) at the chromosomal loci. In both cases, the first intron was removed to exclude potential confounding effects. The *rap1*^+^ gene carrying the *sgo2*^+^ intron (*rap1^sgo2-intron^*) exhibited a marked reduction in Rap1 protein levels (Supplementary Figure S2), suggesting that both intronic sequences and their combination with flanking exons may contribute to proper Rap1 expression. In contrast, the *poz1*^+^ gene containing the *sgo2*^+^ intron (*poz1^sgo2-intron^*) produced Poz1 protein at levels comparable to those of the wild-type and first intron-deleted *poz1*^+^ strains (*poz1^wt^* and *poz1^poz1-intron^*, respectively), indicating that the second intron of *poz1*^+^ can be functionally replaced by the fourth intron of *sgo2*^+^ (Figure 4C, the WT lanes).

To determine whether the inserted *sgo2*^+^ intron in *poz1*^+^ is subject to regulation by Tls1 and Cay1, we deleted *tls1*^+^ and *cay1*^+^ in the *poz1^sgo2-intron^* background. Neither deletion affected Poz1 protein levels in this strain, whereas significant reductions were observed in the *poz1^poz1-intron^* strain upon deletion of *tls1*^+^ or *cay1*^+^ (Figure 4C). These findings indicate that the *sgo2*^+^ intron in *poz1*^+^ is not regulated by Tls1 or Cay1 but likely depends on other splicing factors. Collectively, these results suggest that the intact intron sequence of *poz1*^+^ is required for the specific regulation by Tls1 and Cay1.

### Tls1 and Cay1 exhibit distinct modes of association with the splicing machinery

To investigate how Tls1 and Cay1 contribute to the splicing of *rap1*^+^ and *poz1*^+^ transcripts, we first tested whether either protein exhibits specific affinity for their intronic sequences. We performed *in vitro* RNA pull-down assays by incubating *S. pombe* whole cell extracts with GST-tagged Tls1 or Cay1 proteins purified from *E. coli* (Figure 5A). RNAs bound to the GST proteins were then isolated and analyzed by RT-qPCR. The introns of *rap1*^+^ and *poz1*^+^ were significantly enriched in the GST-Cay1 samples compared to the intron of *sgo2*^+^ or the corresponding exonic regions. In contrast, GST-Tls1 did not show substantial enrichment of any RNA (Figure 5B). These results suggest that Cay1, but not Tls1, may participate in the splicing of *rap1*^+^ and *poz1*^+^ transcripts through direct interaction with their introns.

**Figure 5.**
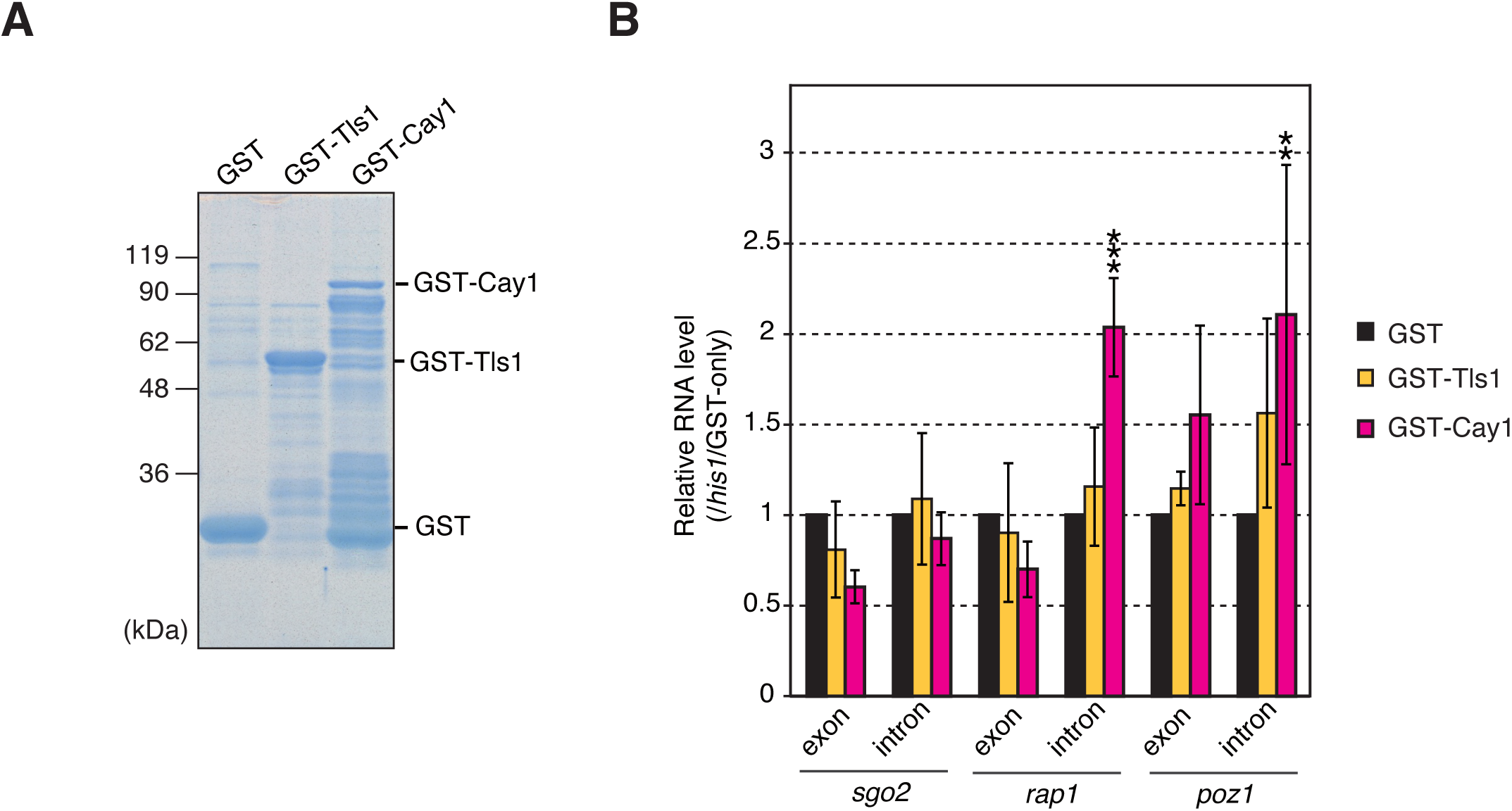
Cay1 associates with the introns of *rap1*^+^ and *poz1*^+^ *in vitro*. (A) Purification of GST, GST-Tls1, and GST-Cay1 proteins from *E. coli*. Proteins bound to glutathione beads were analyzed by SDS-PAGE followed by CBB staining. (B) *In vitro* RNA-protein binding assay. Whole-cell extracts from the wild-type strain were incubated with GST, GST-Tls1, or GST-Cay1, and the bound RNAs were analyzed by RT-qPCR. Each value was normalized first to *his1*^+^ (exon) and then to the GST control. Error bars represent the s.d. (*N* = 3 biologically independent experiments). Student’s t-test was performed against GST. \*\*\**p* ≤ 0.001; 0.001 < \*\**p* ≤ 0.01.

We next examined whether Tls1 and Cay1 interact with spliceosomal components. The spliceosome, which catalyzes pre-mRNA splicing, consists of small nuclear ribonucleoprotein (snRNP) complexes—U1, U2, U4, U5, and U6—as well as non-snRNP complexes such as the NineTeen Complex (NTC) (19–21) (Figure 6A). Co-immunoprecipitation using anti-Tls1 antibodies was unsuccessful due to the instability of Tls1 in *S. pombe* cell extracts. Therefore, we again employed *in vitro* pull-down assays using GST-Tls1 or GST-Cay1 proteins.

**Figure 6.**
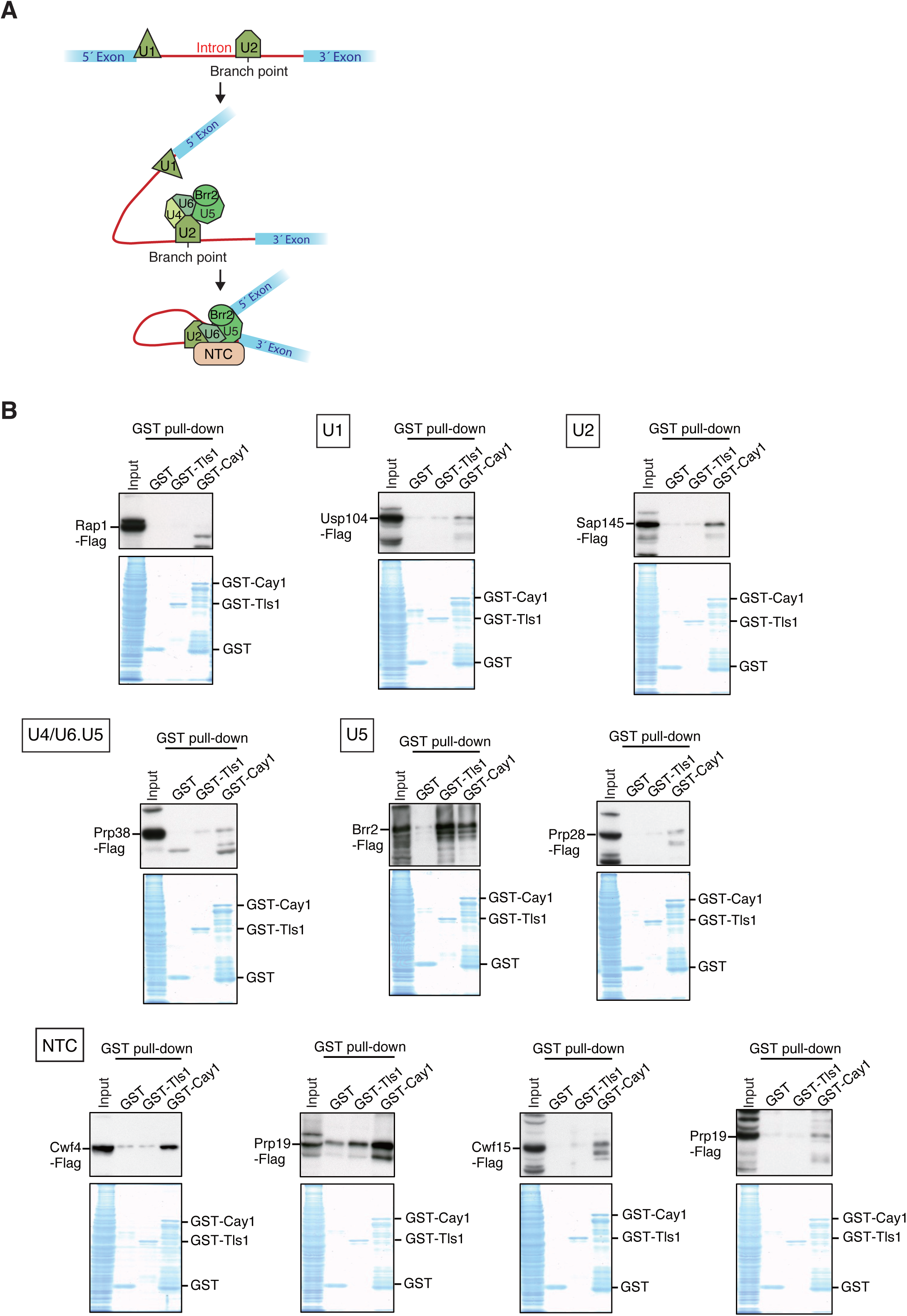
Tls1 and Cay1 associate with spliceosome subunits. (A) Schematic illustration of the spliceosome during pre-mRNA splicing. (B) GST pull-down assay. Whole-cell extracts from strains expressing each Flag-tagged protein were incubated with GST, GST-Tls1, or GST-Cay1, and the bound proteins were analyzed by western blotting with anti-Flag antibody (upper). The subunits to which the proteins belong are indicated in squares. The same samples in each assay were also analyzed by SDS-PAGE and CBB staining (lower).

Consistent with a previous report (13), GST-Tls1 exhibited strong affinity for Brr2 among the spliceosomal proteins tested (Figure 6B). Brr2 is a DEAD/DEAH-box ATPase component of the U5 snRNP that promotes U4/U6 dissociation during spliceosome activation (22–25). Notably, Tls1, but not Cay1, directly interacted with Brr2 in a yeast two-hybrid assay, supporting the possibility of a direct protein-protein interaction (Supplementary Figure S3). GST-Tls1 also showed weaker binding to Prp19, a subunit of the NTC, but not to other spliceosomal components or Rap1 (Figure 6B). In contrast, GST-Cay1 displayed interactions with multiple spliceosome proteins, particularly those associated with U5 and the NTC (Figure 6B). These findings suggest that Tls1 and Cay1 associate with the spliceosome through distinct sets of interactions, likely reflecting their separate roles in splicing regulation.

### Abnormal telomere elongation caused by impaired splicing of *rap1*^+^ and *poz1*^+^

We next investigated the physiological significance of RNA splicing regulation of *rap1*^+^ and *poz1*^+^ by Tls1 and Cay1. Because Rap1 and Poz1 are key regulators of telomere length (3,6,7), we analyzed telomere length in various mutant strains. The *cay1*Δ mutant exhibited substantially longer telomeres than the *tls1*Δ mutant, and the *tls1*Δ*cay1*Δ double mutant showed even greater telomere elongation than *cay1*Δ alone (Figure 7A). These findings correlate with the synergistic effects of Tls1 and Cay1 deletions on Rap1 and Poz1 protein levels (Figure 2). Notably, no additional telomere elongation was observed in *rap1*Δ*poz1*Δ cells upon deletion of Tls1 and Cay1, suggesting that Tls1 and Cay1 regulate telomere length predominantly through Rap1 and Poz1, rather than through other factors (Figure 7A).

**Figure 7.**
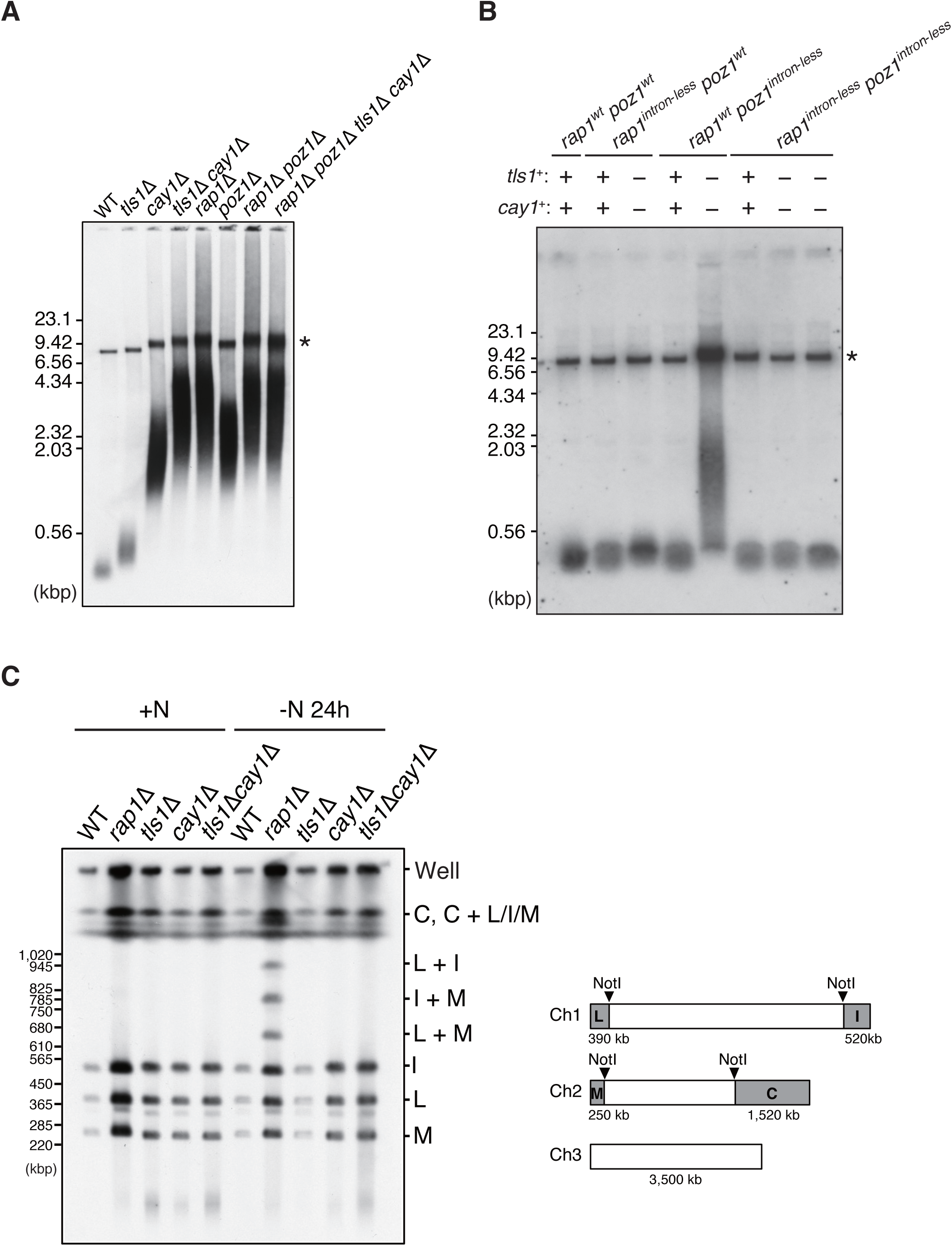
Effects of Tls1 and Cay1 deletion on telomere length and chromosome-end structure. (A) Analysis of telomere length in various strains. Genomic DNA was digested with ApaI, separated in a 1% agarose gel, and analyzed by Southern blotting using telomeric DNA as a probe. An asterisk indicates the position of chromosome 3 telomeres. (B) Analysis of telomere length in strains expressing *rap1^wt^poz1^wt^*, *rap1^intron-less^poz1^wt^*, *rap1^wt^poz1^intron-less^*, or *rap1^intron-less^poz1^intron-less^*. Each genetic background for *tls1*^+^ and *cay1*^+^ is indicated by + or –. Genomic DNA was digested with ApaI, separated in a 1% agarose gel, and analyzed by Southern blotting using telomeric DNA as a probe. An asterisk indicates the position of chromosome 3 telomeres. (C) Analysis of chromosome end structure in various strains. Cells were grown in nitrogen-containing medium (+N) or transferred to nitrogen-free medium and cultured for 24 h (-N). Genomic DNA was digested with NotI and analyzed by PFGE followed by Southern blotting using telomeric DNA as a probe (left). The right panel illustrates the telomere-containing NotI restriction fragments, with approximate lengths indicated below each box. Note that chromosome 3, which contains no NotI site, was mostly retained in the wells of the gel due to its long length.

To determine whether the abnormal telomere elongation in *tls1*Δ*cay1*Δ cells results from defective splicing of *rap1*^+^ and *poz1*^+^ transcripts, we analyzed telomere lengths in strains with chromosomal deletions of the introns of these genes. Deletion of all introns from *rap1*^+^, *poz1*^+^, or both genes had no apparent effect on telomere length compared with wild-type cells (Figure 7B), indicating that the introns themselves are dispensable for telomere length control under normal conditions. In *tls1*Δ*cay1*Δ cells lacking either *rap1*^+^ or *poz1*^+^ introns, slight or moderate telomere elongation was observed, respectively, likely reflecting residual splicing defects of *poz1*^+^ or *rap1*^+^ transcripts (Figure 7B). In contrast, *tls1*Δ*cay1*Δ cells lacking introns in both *rap1*^+^ and *poz1*^+^ exhibited telomere lengths comparable to those in wild-type cells (Figure 7B), indicating that telomere elongation phenotype in *tls1*Δ*cay1*Δ cells is entirely attributable to defective splicing of *rap1*^+^ and *poz1*^+^ transcripts. These results suggest that telomere length is highly sensitive to reductions in Rap1 and Poz1 protein levels, particularly to the amount of Rap1.

### Deletion of Tls1 and Cay1 does not impair chromosome end protection despite reduced Rap1 levels

It has been shown that Rap1 is required for chromosome end protection in G_1_ phase, when the non-homologous end joining (NHEJ) pathway is active (8,26,27). Accordingly, chromosome end fusions occur in *rap1*Δ cells arrested in G_1_ phase under nitrogen starvation (Figure 7C). Notably, the *tls1*Δ*cay1*Δ mutant, as well as the *cay1*Δ and *tls1*Δ single mutants, in which Rap1 protein levels are substantially reduced, exhibited no detectable chromosome end fusions in G_1_ (Figure 7C). These findings indicate that a very small amount of Rap1 is sufficient to suppress NHEJ at chromosome ends. Thus, in contrast to telomere length, which is highly sensitive to reductions in Rap1 levels, chromosome end protection remains fully functional even with markedly reduced amounts of Rap1.

### Defective splicing of *rap1*^+^ partially accounts for abnormal spore formation in ***tls1*Δ*cay1*Δ cells**

Rap1 also plays a crucial role in meiotic chromosome dynamics in *S. pombe* (6,7). Upon nitrogen starvation, *S. pombe* cells arrest in G_1_ phase, mate with cells of the opposite mating type, and subsequently enter meiosis. After completion of the first and second meiotic divisions, spores are formed around the segregated chromosome sets. Prior to mating, telomeres are clustered at the spindle pole body (SPB) through interactions among Taz1, Rap1, the inner nuclear membrane proteins Bqt3 and Bqt4, and meiosis-specific proteins Bqt1 and Bqt2 (8,9,28) (Figure 1A). Following mating, the SPB-telomere complex drives dynamic chromosome movements, known as “horsetail movement,” characterized by figure-eight-shaped nuclear oscillations that promote homologous chromosome pairing for meiotic recombination (29,30). In addition, the SPB-telomere association is required for proper meiotic chromosome segregation. Consequently, defective meiosis and abnormal spore formation are observed in *rap1*Δ cells (6–8). Consistent with previous reports (6–8), the *rap1*Δ mutant frequently formed abnormal numbers (one to three) of spores, whereas almost all wild-type and *poz1*Δ strains produced four spores under nitrogen starvation (Figure 8A and B). Furthermore, the *rap1*Δ*poz1*Δ double mutant exhibited sporulation defects comparable to those of the *rap1*Δ mutant (Figure 8B), indicating that Poz1 is dispensable for normal spore formation.

**Figure 8.**
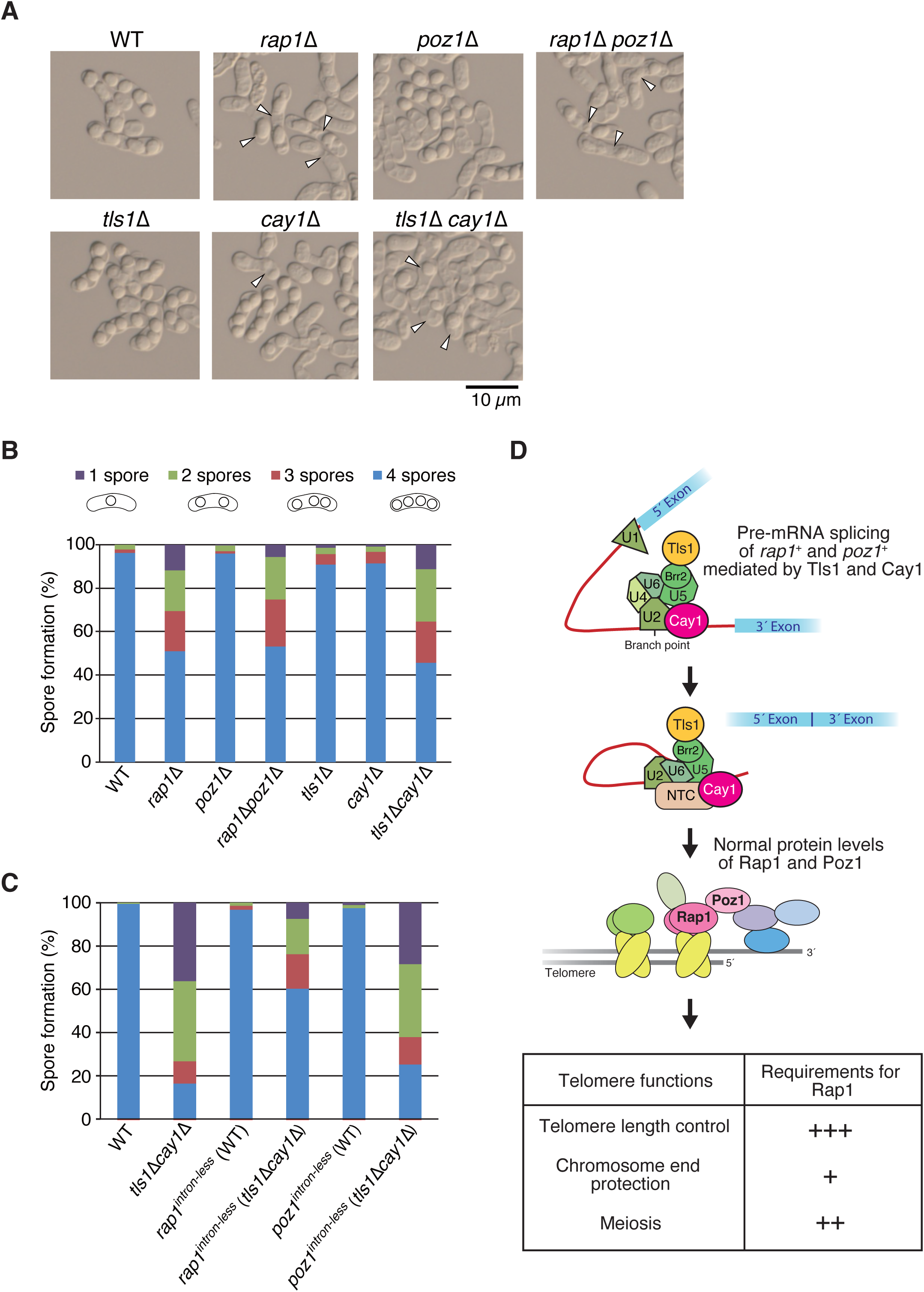
Effects of Tls1 and Cay1 deletion on spore formation. (A) Spore formation in various strains. Homothallic haploid cells were grown on MEA medium at 28°C for 2 days to induce mating, meiosis, and spore formation. DIC images are shown. Arrowheads indicate abnormal spore formation. (B) Frequency of abnormal spore formation in each strain under the same conditions as in (A). More than 200 asci were analyzed for each strain. (C) Frequency of abnormal spore formation in each strain, as in (B). (D) Model of the involvement of Tls1 and Cay1 in pre-mRNA splicing and protein expression of *rap1*^+^ and *poz1*^+^. Tls1 shows a strong affinity for Brr2, whereas Cay1 exhibits relatively strong affinities for *rap1*^+^ and *poz1*^+^ introns, as well as for U5 and NTC. Efficient splicing mediated by Tls1 and Cay1 ensures normal protein levels of Rap1 and Poz1 (upper). Different levels of requirement for Rap1 in the three telomere functions are indicated (lower).

The *tls1*Δ and *cay1*Δ single mutants, which exhibit reduced Rap1 protein levels, showed only mild sporulation defects compared with the *rap1*Δ mutant. In contrast, the *tls1Δcay1Δ* double mutant, which shows further reduction of Rap1 protein levels compared with each single mutant, displayed severe sporulation defects similar to those of *rap1*Δ cells (Figure 8A and B). These results suggest that abnormal spore formation in *tls1Δcay1Δ* cells is at least partly attributable to reduced Rap1 protein levels.

To further assess whether these defects arise from defective splicing of *rap1*^+^ or *poz1*^+^ transcripts, we examined sporulation in strains lacking introns of each gene. Elimination of all introns from *rap1*^+^ or *poz1*^+^ had no impact on spore formation; these strains produced spores normally, as did wild-type cells (Figure 8C). In contrast, *tls1Δcay1Δ* cells lacking *poz1*^+^ introns showed severe sporulation defects, whereas *tls1Δcay1Δ* cells lacking *rap1*^+^introns exhibited only moderate defects, less severe than those in *tls1Δcay1Δ* cells with intact *rap1*^+^ introns (Figure 8C). These results indicate that abnormal spore formation in *tls1Δcay1Δ* cells is partially due to defective splicing of *rap1*^+^ transcripts. However, the persistence of sporulation defects even in the absence of *rap1*^+^ introns suggests that Tls1 and Cay1 also regulate the pre-mRNA splicing of other genes important for normal meiosis. Collectively, these findings suggest that a relatively small amount of Rap1 is sufficient to support normal meiosis and spore formation. Furthermore, meiosis is less sensitive to reductions of Rap1 protein levels than telomere length control, but more sensitive than chromosome end protection.

## Discussion

In this study, we demonstrate that efficient pre-mRNA splicing of *rap1*^+^ and *poz1*^+^ mediated by Tls1 and Cay1 is essential for maintaining adequate levels of Rap1 and Poz1 proteins. Tls1 strongly associates with Brr2, whereas Cay1 preferentially associates with introns as well as multiple spliceosome subunits. Importantly, we reveal that distinct telomere functions exhibit differential quantitative requirements for Rap1: telomere length regulation and, to a lesser extent, meiosis require higher protein levels, whereas chromosome end protection can be sustained with minimal amounts. These findings establish a hierarchical framework for understanding how splicing-dependent regulation of shelterin components coordinates multiple aspects of telomere biology (Figure 8D).

In *tls1*Δ cells, we observed only moderate telomere DNA elongation (∼400 bp) compared with the wild type (∼300 bp), which is substantially smaller than previously reported (∼2 kb longer than wild type) (13). Multiple independently generated *tls1*Δ strains confirmed this modest effect (Figure 1B), indicating that Tls1 exerts a relatively minor influence on telomere length.

Notably, in single (*tls1*Δ or *cay1*Δ) and double (*tls1*Δ*cay1*Δ) deletion mutants, splicing of *rap1*⁺ and *poz1*⁺ transcripts is not completely abolished; the effect on *poz1*⁺ splicing, in particular, is mild even in the double mutant (Figure 3C and D). This indicates that Tls1 and Cay1 do not serve as absolutely essential factors for splicing but rather function to promote efficient pre-mRNA processing of *rap1*⁺ and *poz1*⁺. Interestingly, while the impact of Tls1 or Cay1 deletion on splicing itself is relatively mild, the protein levels of Rap1 and Poz1 are drastically reduced (Figure 2). This could suggest that Tls1 and Cay1 might have additional, splicing-independent roles in maintaining protein levels. However, when all introns were removed from *rap1*⁺ and *poz1*⁺, the double deletion of Tls1 and Cay1 no longer caused a reduction in protein levels (Figure 4A and B). This confirms that defective splicing is indeed the primary cause of the decreased Rap1 and Poz1 protein abundance. A remaining question is why, in Tls1 and Cay1 deletion mutants, substantial amounts of correctly spliced *rap1*⁺ and *poz1*⁺ RNA are still present, yet protein levels are severely reduced. At present, the mechanism is unclear. One possibility is that Tls1 and Cay1 have important roles immediately after splicing, perhaps facilitating efficient translation or protein stability. Alternatively, reduced expression of other target genes regulated by Tls1 and Cay1 may indirectly exacerbate the decrease in Rap1 and Poz1 protein levels.

Analysis of single and double *tls1*Δ and *cay1*Δ mutants further revealed the differential Rap1 requirements for specific telomere functions. For telomere length regulation, a sufficient amount of shelterin proteins must bind telomere ends to enable efficient interactions between double-stranded and single-stranded telomeric DNA via the shelterin complex; insufficient protein levels lead to defects in length control. In contrast, chromosome end protection requires only minimal amounts of Rap1 to prevent chromosome end fusions. Meiosis shows an intermediate requirement: single mutants display little or no defect, whereas the double mutant exhibits severe abnormalities (Figure 8C). This indicates that the Rap1 levels in single mutants are sufficient for meiosis, but those in the double mutant fall below a critical threshold. During meiotic prophase, telomeres cluster at the SPB and engage in horsetail movement to facilitate homolog pairing; an adequate amount of Rap1 is likely required to maintain this dynamic movement. Collectively, our findings reveal a hierarchy of Rap1 requirements across telomere functions and highlight splicing-dependent regulation as a key mechanism coordinating multiple aspects of telomere biology.

## Acknowledgments

We thank A. Koyanagi, M. Tanaka, Y. Takeshita, and F. Ishikawa for their supports of initial stage of this study, and all former and present lab members for discussion and support.

## Author Contributions

J.K. conceived the project. J.K. and M.T. designed and performed experiments and analyzed data. J.K. and Y.O. wrote the manuscript.

## Funding

Japan Society for the Promotion of Science (JSPS) KAKENHI [JP17H03606, JP21H00244, JP22H04685, JP23H02408], Daiichi Sankyo Foundation of Life Science, Ohsumi Frontier Science Foundation, and Bioscience Research Grant of Takeda Science Foundation (to J.K.).

## Competing interests

The authors declare no competing interests.

## Notes

### Competing Interest Statement

The authors have declared no competing interest.

