## Supplementary Data for "Differential quantitative requirements for pre-mRNA splicing-regulated shelterin protein levels in distinct telomere functions"

**for**

Contents: Figures S1–S3

Tables S1–S2

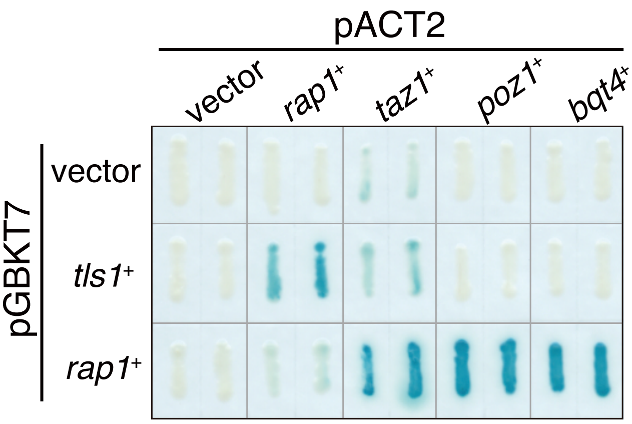


**Figure S1. Rip6/Tls1 was identified as a Rap1-interacting protein in an *S. cerevisiae* two-hybrid system.**

The *S. cerevisiae* Y190 strain was co-transformed with two plasmids, pACT2 and pGBKT7, carrying various telomere-related genes. The vectors carrying no gene were used as a negative control. Each transformant was grown on selective medium and assayed for β-galactosidase activity. Transformants with pGBKT7-*rap1*$^{+}$ and pACT2-*taz1*$^{+}$, *poz1*$^{+}$ or *bqt4*$^{+}$ served as positive controls.

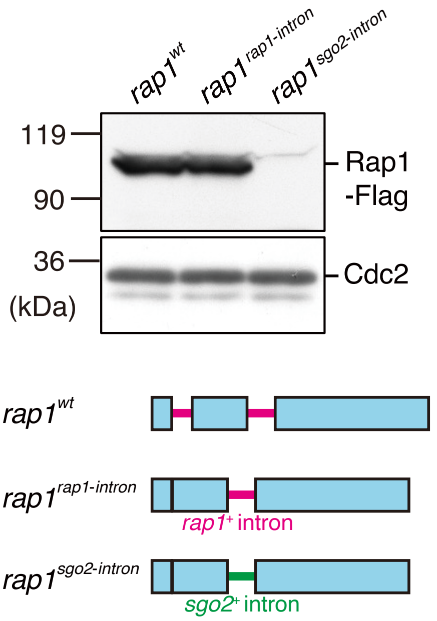


**Figure S2. Marked decrease in Rap1 protein levels when the *rap1*$^+$ intron was replaced with the *sgo2*$^+$ intron.**

Whole-cell extracts from strains expressing *rap1*$^{wt}$, *rap*$^{rap1\text{-}intron}$, or *rap1*$^{sgo2\text{-}intron}$ were analyzed by western blotting with anti-Flag (Rap1) or anti-PSTAIRE (for Cdc2, loading control) antibodies.

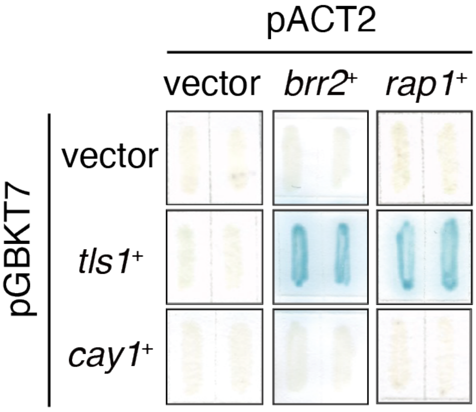


**Figure S3. Tls1 associates with Brr2 in an *S. cerevisiae* two-hybrid system.**

The *S. cerevisiae* Y190 strain was co-transformed with two plasmids as described in Figure S1. Each transformant was grown on selective medium and assayed for β-galactosidase activity. Transformants with pGBKT7-*tls1*$^{+}$ and pACT2-*rap1*$^{+}$ served as a positive control.

### Table S1. *S. pombe* strains used in this study

**Fig. 1B**

| | |
|---|---|
| JK107 | $h^-$ |
| JP2724 | $h^-$ *tls1::hphMX6* |
| RT4413 | $h^-$ *tls1::natMX6* |
| JP2962 | $h^-$ *leu1-32 ura4-D18* |
| JP2480 | $h^-$ *leu1-32 ura4-D18 tls1::ura4* |
| RT4461 | $h^-$ *leu1-32 ura4-D18 tls1::ura4* |

**Fig. 2**

| | |
|---|---|
| JK107 | $h^-$ |
| RT4413 | $h^-$ *tls1::natMX6* |
| JP3683 | $h^-$ *cay1::kanMX6* |
| RT4415 | $h^-$ *tls1::natMX6 cay1::kanMX6* |
| JP3228 | $h^-$ *poz1-3flag-kanMX6* |
| JP3230 | $h^-$ *poz1-3flag-kanMX6 tls1::hphMX6* |
| JP3904 | $h^-$ *poz1-3flag-hphMX6 cay1::kanMX6* |
| JP3933 | $h^-$ *poz1-3flag-hphMX6 tls1::hphMX6 cay1::kanMX6* |
| JP2810 | $h^-$ *taz1-3flag-kanMX6* |
| JP3232 | $h^-$ *taz1-3flag-kanMX6 tls1::hphMX6* |
| JP3940 | $h^-$ *taz1-3flag-kanMX6 cay1::hphMX6* |
| RT4547 | $h^-$ *taz1-3flag-kanMX6 tls1::natMX6 cay1::hphMX6* |
| RT5295 | $h^-$ *pot1-3flag-hphMX6* |
| RT5297 | $h^-$ *pot1-3flag-hphMX6 tls1::natMX6* |
| RT5299 | $h^-$ *pot1-3flag-hphMX6 cay1::kanMX6* |
| RT5301 | $h^-$ *pot1-3flag-hphMX6 tls1::natMX6 cay1::kanMX6* |
| RT5287 | $h^-$ *tpz1-3flag-hphMX6* |
| RT5289 | $h^-$ *tpz1-3flag-hphMX6 tls1::natMX6* |
| RT5291 | $h^-$ *tpz1-3flag-hphMX6 cay1::kanMX6* |
| RT5293 | $h^-$ *tpz1-3flag-hphMX6 tls1::natMX6 cay1::kanMX6* |
| RT5279 | $h^-$ *ccq11-3flag-hphMX6* |
| RT5281 | $h^-$ *ccq1-3flag-hphMX6 tls1::natMX6* |
| RT5283 | $h^-$ *ccq1-3flag-hphMX6 cay1::kanMX6* |
| RT5285 | $h^-$ *ccq1-3flag-hphMX6 tls1::natMX6 cay1::kanMX6* |
| JP3224 | $h^-$ *rif1-3flag-kanMX6* |

| | |
|---|---|
| JP3226 | $h^-$ *rif1-3flag-kanMX6 tls1::hphMX6* |
| JP3942 | $h^-$ *rif1-3flag-kanMX6 cay1::hphMX6* |
| RT4549 | $h^-$ *rif1-3flag-kanMX6 tls1::natMX6 cay1::hphMX6* |
| RT5456 | $h^-$ *sgo2-3flag-hphMX6* |
| RT5461 | $h^-$ *sgo2-3flag-hphMX6 tls1::natMX6* |
| RT5463 | $h^-$ *sgo2-3flag-hphMX6 cay1::kanMX6* |
| RT5465 | $h^-$ *sgo2-3flag-hphMX6 tls1::natMX6 cay1::kanMX6* |

**Fig. 3B, C, D**

| | |
|---|---|
| JK107 | $h^-$ |
| RT4413 | $h^-$ *tls1::natMX6* |
| JP3683 | $h^-$ *cay1::kanMX6* |
| RT4415 | $h^-$ *tls1::natMX6 cay1::kanMX6* |

**Fig. 4A**

| | |
|---|---|
| JP2983 | $h^-$ *rap1-3flag-hphMX6* |
| RT5431 | $h^-$ *rap1*$^{intron\text{-}less}$*-3flag-hphMX6* |
| RT5433 | $h^-$ *rap1*$^{intron\text{-}less}$*-3flag-hphMX6 tls1::natMX6* |
| RT5435 | $h^-$ *rap1*$^{intron\text{-}less}$*-3flag-hphMX6 cay1::kanMX6* |
| RT5443 | $h^-$ *rap1*$^{intron\text{-}less}$*-3flag-hphMX6 tls1::natMX6 cay1::kanMX6* |

**Fig. 4B**

| | |
|---|---|
| JP3228 | $h^-$ *poz1-3flag-kanMX6* |
| RT5432 | $h^-$ *poz1*$^{intron\text{-}less}$*-3flag-hphMX6* |
| RT5439 | $h^-$ *poz1*$^{intron\text{-}less}$*-3flag-hphMX6 tls1::natMX6* |
| RT5441 | $h^-$ *poz1*$^{intron\text{-}less}$*-3flag-hphMX6 cay1::kanMX6* |
| RT5444 | $h^-$ *poz1*$^{intron\text{-}less}$*-3flag-hphMX6 tls1::natMX6 cay1::kanMX6* |

**Fig. 4C**

| | |
|---|---|
| JP3228 | $h^-$ *poz1-3flag-kanMX6* |
| RT5469 | $h^-$ *poz1*$^{poz1\text{-}intron}$*-3flag-hphMX6* |
| RT5484 | $h^-$ *poz1*$^{poz1\text{-}intron}$ *-3flag-hphMX6 tls1::natMX6* |
| RT5486 | $h^-$ *poz1*$^{poz1\text{-}intron}$ *-3flag-hphMX6 cay1::kanMX6* |
| RT5470 | $h^-$ *poz1*$^{sgo2\text{-}intron}$*-3flag-hphMX6* |
| RT5488 | $h^-$ *poz1*$^{sgo2\text{-}intron}$ *-3flag-hphMX6 tls1::natMX6* |
| RT5490 | $h^-$ *poz1*$^{sgo2\text{-}intron}$ *-3flag-hphMX6 cay1::kanMX6* |

**Fig. 5B**

| | |
|---|---|
| JK107 | $h^-$ |

**Fig. 6B**

| | |
|---|---|
| JP2983 | $h^-$ *rap1-3flag-hphMX6* |
| RT4775 | $h^-$ *usp104-3flag-kanMX6* |
| RT4787 | $h^-$ *sap145-3flag-kanMX6* |
| RT4791 | $h^-$ *prp38-3flag-kanMX6* |
| RT4453 | $h^-$ *brr2-3flag-kanMX6* |
| RT4551 | $h^-$ *prp28-3flag-kanMX6* |
| RT4553 | $h^-$ *cwf4-3flag-kanMX6* |
| RT4555 | $h^-$ *prp19-3flag-kanMX6* |
| RT4783 | $h^-$ *cwf15-3flag-kanMX6* |
| RT4555 | $h^-$ *prp19-3flag-kanMX6* |

**Fig. 7A**

| | |
|---|---|
| JK107 | $h^-$ |
| RT4413 | $h^-$ *tls1::natMX6* |
| JP3683 | $h^-$ *cay1::kanMX6* |
| RT4415 | $h^-$ *tls1::natMX6 cay1::kanMX6* |
| JM2518 | $h^-$ *rap1::kanMX6* |
| JP3352 | $h^-$ *poz1::hphMX6* |
| RT4417 | $h^-$ *rap1::kanMX6 poz1::hphMX6* |
| RT4421 | $h^-$ *rap1::kanMX6 poz1::natMX6 tls1::hphMX6 cay1::BSD* |

**Fig. 7B**

| | |
|---|---|
| JK107 | $h^-$ |
| RT5431 | $h^-$ *rap1*$^{intron\text{-}less}$*-3flag-hphMX6* |
| RT5443 | $h^-$ *rap1*$^{intron\text{-}less}$*-3flag-hphMX6 tls1::natMX6 cay1::kanMX6* |
| RT5432 | $h^-$ *poz1*$^{intron\text{-}less}$*-3flag-hphMX6* |
| RT5444 | $h^-$ *poz1*$^{intron\text{-}less}$*-3flag-hphMX6 tls1::natMX6 cay1::kanMX6* |
| JP5540 | $h^-$ *rap1*$^{intron\text{-}less}$*-3flag-hphMX6 poz1*$^{intron\text{-}less}$*-3flag-hphMX6* |
| JP5547 | $h^-$ *rap1*$^{intron\text{-}less}$*-3flag-hphMX6 poz1*$^{intron\text{-}less}$*-3flag-hphMX6* *tls1::natMX6 cay1::kanMX6* |

**Fig. 7C**

| | |
|---|---|
| JK107 | $h^{-}$ |
| JM2518 | $h^{-}$ *rap1*::*kanMX6* |
| RT4413 | $h^{-}$ *tls1*::*natMX6* |
| JP3683 | $h^{-}$ *cay1*::*kanMX6* |
| RT4415 | $h^{-}$ *tls1*::*natMX6 cay1*::*kanMX6* |

**Fig. 8A, B**

| | |
|---|---|
| JP245 | $h^{90}$ *ade6-M210 leu1-32 ura4-D18* |
| RT5364 | $h^{90}$ *ade6-M210 leu1-32 ura4-D18 rap1*::*kanMX6* |
| HI4283 | $h^{90}$ *ade6-M210 leu1-32 ura4-D18 poz1*::*ura4* |
| RT5366 | $h^{90}$ *ade6-M210 leu1-32 ura4-D18 rap1*::*kanMX6 poz1*::*ura4* |
| JP2482 | $h^{90}$ *ade6-M210 leu1-32 ura4-D18 tls1*::*ura4* |
| RT4467 | $h^{90}$ *ade6-M210 leu1-32 ura4-D18 cay1*::*ura4* |
| RT4541 | $h^{90}$ *ade6-M210 leu1-32 ura4-D18 tls1*::*ura4 cay1*::*hphMX6* |

**Fig. 8C**

| | |
|---|---|
| JP245 | $h^{90}$ *ade6-M210 leu1-32 ura4-D18* |
| RT4541 | $h^{90}$ *ade6-M210 leu1-32 ura4-D18 tls1*::*ura4 cay1*::*hphMX6* |
| JP5500 | $h^{90}$ *ade6-M210 leu1-32 ura4-D18* $rap1^{intron\text{-}less}$*-3flag-hphMX6* |
| JP5517 | $h^{90}$ *ade6-M210 leu1-32 ura4-D18* $rap1^{intron\text{-}less}$*-3flag-hphMX6* *tls1*::*natMX6 cay1*::*kanMX6* |
| JP5502 | $h^{90}$ *ade6-M210 leu1-32 ura4-D18* $poz1^{intron\text{-}less}$*-3flag-hphMX6* |
| JP5521 | $h^{90}$ *ade6-M210 leu1-32 ura4-D18* $poz1^{intron\text{-}less}$*-3flag-hphMX6* *tls1*::*natMX6 cay1*::*kanMX6* |

**Fig. S2**

| | |
|---|---|
| JP2983 | $h^{-}$ *rap1-3flag-hphMX6* |
| RT5454 | *h-* $rap1^{rap1\text{-}intron}$*-3flag-hphMX6* |
| RT5471 | *h-* $rap1^{sgo2\text{-}intron}$*-3flag-hphMX6* |

### Table S2. Primers used in this study

***his1***$^{+}$**-exon**

jk1336 5′-TGTCCACCTCGGAATCACTG-3′

jk1335 5′-CGAAGACGTGCTTCAGCGA-3′

***sgo2***$^{+}$**-exon**

jk1711 5′-CACGTAGCTCCTTTGCTACCAAC-3′

rt250 5′-CGTTCTCATTAGGGGATTTCAAA-3′

***sgo2***$^{+}$**-intron**

rt248 5′-GAAGACCTTGACGAGGTAAGTTGAAT-3′

rt249 5′-CACGACGGATTCCTTTCAGAC-3′

***rap1***$^{+}$**-exon**

jk1599 5′-AAATCACCGTGAGGTTGATGAAG-3′

jk1600 5′-ACCCGTAGATTCCATAGCTTCAAG-3′

***rap1***$^{+}$**-intron**

rt244 5′-AACTCGCCAGAAAGGTATGATCTT-3′

rt245 5′-CAAGAGAATGTTGAGGATACTAGGAAAC-3′

***poz1***$^{+}$**-exon**

rt148 5′-GATGGAAGCTTTACAACGTTTATC-3′

rt149 5′-CTGAGCACACTATTGAATTTAGAC-3′

***poz1***$^{+}$**-intron**

rt246 5′-GCCGTTGGGTAAGGACCAA-3′

rt247 5′-CTAGAAAATGAAAGGCCATCTAGAATACAT-3′

***rap1***$^{+}$ (for detection of splicing patterns)

rt167 5′-CATATTAGGGGATTCAAAAATGCAATTG-3′

rt168 5′-GATAGAGTTTGTTCATGTGTGCTTTG-3′

***poz1***$^{+}$ (for detection of splicing patterns)

rt169 5′-GAGAAGAATCTTCCATTGTGAACG-3′

rt149 5′-CTGAGCACACTATTGAATTTAGAC-3′
